# Deep Learning-Driven Gender Classification of *Drosophila melanogaster*: Advancing Real-Time Analysis with Mobile Integration

**DOI:** 10.1101/2025.04.04.647141

**Authors:** Jayant Rai, S Aswini, Soumyadeep Paul, Triveni Shelke, Ishaan Gupta

**Author notes:** Correspondence to Ishaan Gupta.

## Abstract

Researchers extensively use *Drosophila melanogaster* as a model organism in genetic research and developmental studies to better understand human diseases such as cancer and diabetes. For such investigations, it is necessary to identify and classify male and female flies. This study uses deep learning techniques for gender classification of *Drosophila melanogaster* via models YOLOv8, Detectron2, ResNet-50, InceptionV3, and MobileNet. Benchmarking revealed that YOLOv8 achieved a maximum accuracy of 98.0%. The classification and object detection models are integrated into a mobile application, facilitating real-time gender classification. This study demonstrates the possibility of integrating deep learning models with mobile technology while ensuring an efficient way for gender prediction, leading to better reproducibility and generating new opportunities in biological research.

## Introduction

Since the early 1900s, *Drosophila melanogaster* has played a crucial role in the development of genetics. Scientists have modeled fruit flies to learn the basic principles of genetic inheritance, developmental biology, cancer research, and pharmacology [1]. Their genome is highly conserved, with homologs for around 75% of genes associated with human diseases [2]. Therefore, these flies are widely used as model organisms for studying developmental biology and physiological processes like circadian rhythms, neural development, and aging. These arthropods provide the opportunity to investigate both environmental and genetic determinants of Parkinson’s disease, representing a substantial repository of information concerning the mechanisms underlying the progression of pathogenesis, thereby facilitating the advancement of different therapeutic strategies [3]. Models of Alzheimer’s disease in Drosophila can provide novel insights into disease processes [4]. The heart structure of a fly is developmentally homologous to that found in vertebrates, hence providing a powerful model for the study of cardiac diseases, developmental malformations, and functional defects in adults.[5]. Methods of testing developmental toxicity include exposing the fly embryos to small molecules, toxicants, and drugs, and also through transgenics approaches.[6] In this species, adult females are generally larger than males [7]. The females have a pointed or elongated abdomen, while males have a less pronounced shape [8]. The females exhibit alternating dark and light bands throughout the posterior part of their abdomen, segmented into seven visible segments under low-power magnification [9]. In contrast, males have a fused structure with only five segments [10]. Males can be identified by the presence of sex combs, which are clusters of about ten stout black bristles located on the uppermost tarsal segment of their forelegs, while females lack these bristles [8]. During their pupal stage, sex combs help distinguish males [11]. The females possess a pointed ovipositor at the tip of their abdomen, while males have darkly pigmented, circular claspers located ventrally at the tip. In the larval stage, males can be differentiated by a large white mass of testicular tissue found in the lateral fat bodies at the anterior end of the posterior third of their bodies, which is visible through the skin [12] [8].Traditionally, *Drosophila melanogaster* is classified by manually sorting each fly under the microscope based on physical characteristics. This is a labor-intensive process, as examining hundreds or thousands of flies under a microscope requires considerable time and effort, slowing the research progress. Minor morphological differences may result in inconsistent results when examining younger or mutant flies. This approach demands expertise, i.e., skilled workers, to guarantee accurate classifications. Manual classification is challenging, particularly for investigations involving huge populations or several generations. Moreover, there can be inconsistencies in classification among researchers.In this era of artificial intelligence, an attempt to automatically classify *Drosophila melanogaster* using machine learning models to increase the ease of classification has been devised via supervised machine learning models with labeled data of both male and female flies. The model uses the labeled data to detect and characterize the morphological traits and classify the female and male flies without human intervention. This study utilizes models : YOLOv8, Detectron2, ResNet-50, InceptionV3, and MobileNet for better accuracy and efficiency. The main aim is to integrate this framework into a mobile application, providing the researcher with a scalable, reproducible, and real-time gender-detecting mechanism.

## Methodology

### Data Collection

The experiment utilized the Oregon-R strain of *Drosophila melanogaster*. This wild-type strain has been a fundamental part of Drosophila research for many years, thanks to its genetically stable and well-characterized background. Originating from Oregon, USA, it has been widely embraced in laboratories around the globe, providing a standardized model for a variety of biological studies. Oregon-R flies display typical phenotypic traits, including brick-red eyes, yellow-brown body coloration, and fully functional wings, making them an excellent reference for genetic, developmental, and behavioral experiments. Culture the flies in sterile glass vials with food, preferably corn starch, at 22 °C. Followed by knocking them out using diethyl ether at different time points of their life circle. Overdose of ether can result in the death of the flies. Alternatively, carbon dioxide can be used as a safer anesthetizing agent. The images of anesthetized flies were captured using a T60N Series stereo microscope under controlled lighting and varying viewpoints (Fig. 1). The images are organized into a two different datasets were prepared from the images i.e. Classification and object detection datasets. The classification dataset includes separate pictures of male and female flies. Furthermore each dataset was divided into training, validation, and test set. Meanwhile, the detection dataset has randomized group images of male and female flies. These images were annotated using MakeSense.AI for bounding box detection (Fig. 2).

**Fig. 1.**
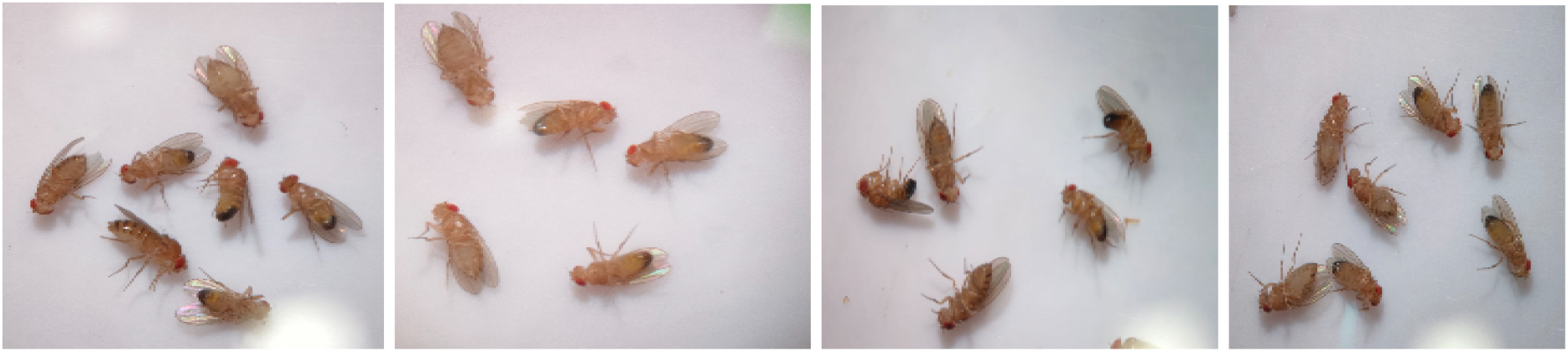
(Data Collection Under Stereo Microscope)

**Fig. 2.**
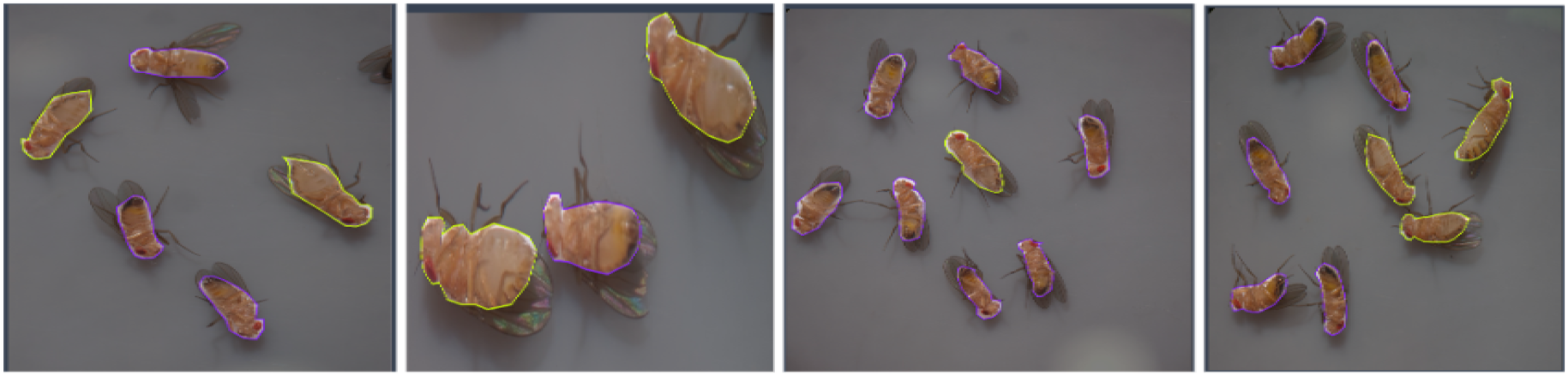
(Annotated Images)

### Data Preprocessing

The data were preprocessed to make the model more reliable and perform well. All images were resized and normalized by the ImageNet mean and standard deviation for the three RGB channels to maintain consistency.

For an efficient learning process, we have split the dataset in an 85:15 ratio for training and validation. To ensure a balanced dataset, we divided the 12,908 images in the classification dataset equally between 6,454 images of males and 6,454 images of females.

We utilized Makesense.ai to label the object detection dataset (Figs. 2, 3). We labeled all the classes in the group’s images to accurately identify the key features for the machine learning model’s performance. We further applied data augmentation methods to improve the diversity and resilience of the training images.

**Fig 3:**
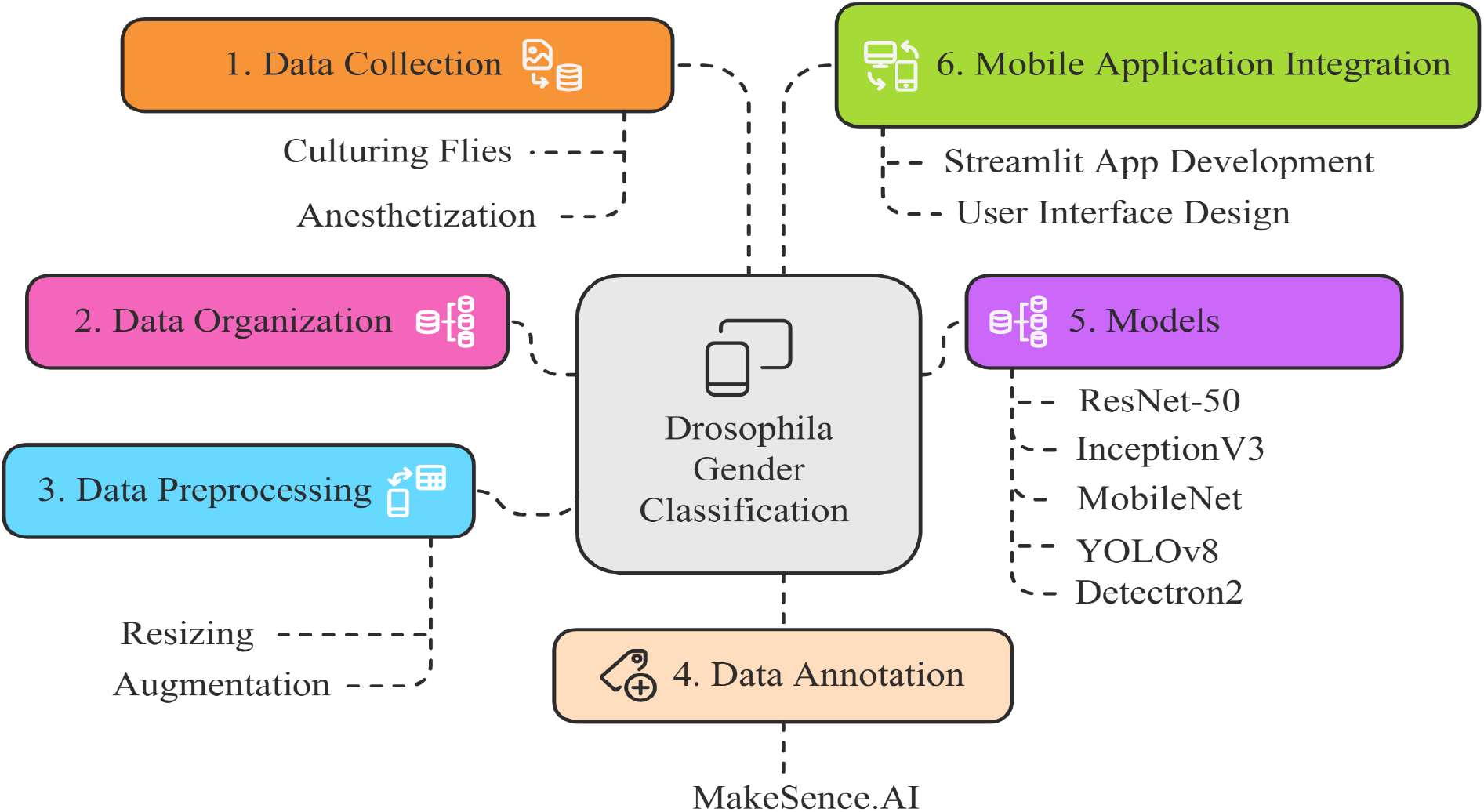
Model Framework. The images of individual flies were subjected to classification after preprocessing using architectures like ResNet -50, InceptionV3, and MobileNet. Images of a group of flies were annotated and subjected to object detection using YOLOv8 and Detectron2 models.

**Fig 4.**
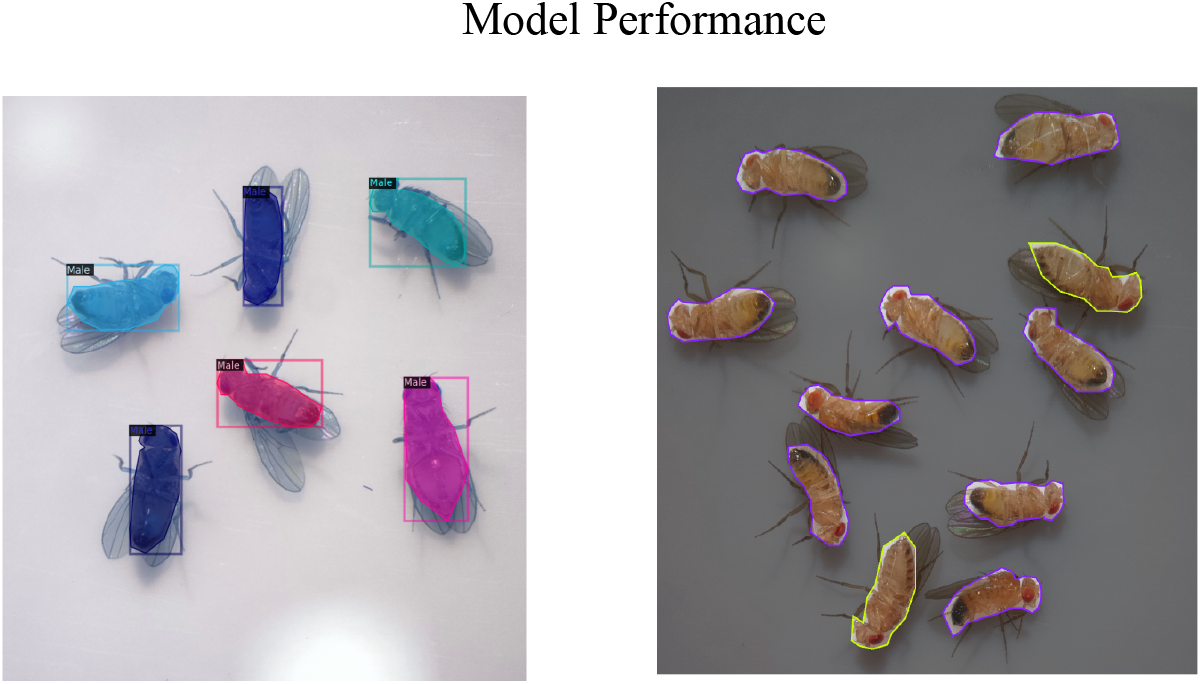
(Results: Detectron2, Yolov8 Respectively)

## Model Architectures

### 1. ResNet-50

ResNet-50 has convolutional layers that extract features from the input image; the identity and convolutional blocks take the features and process and transform them; and the fully connected layers classify the images. ResNet-50 was initially trained on the large ImageNet dataset and achieved a human-level error rate, which allows its use as a powerful model for many image classification tasks. ResNet-50 was pre-trained on ImageNet and used for feature extraction. Layers up to 140 were frozen, and a Global Average Pooling layer was applied to reduce spatial dimensions. A dense softmax classifier was added for binary classification (male/female). [16]

### 2. InceptionV3

Transfer Learning with InceptionV3 benefits from its unique architectural design of Inception blocks, which combine multiple kernel sizes for effective multiscale feature extraction. Complex datasets require more local details which is why the input size is preferably large. It also unearths the last 30 layers to allow the model to learn specialized characteristics of the website domain.[17]

### 3. MobileNet

Chosen for its lightweight architecture suitable for deployment on mobile platforms. Pretrained layers were frozen, followed by fine tuning for classification tasks. Depthwise separable convolutions efficiently separate spatial and channel operations to reduce computational cost. It has a compact design optimized for lightweight deployment with width (α) and resolution (ρ) multipliers for balancing speed and accuracy and suitable for mobile and edge devices. [18]

### 4. YOLOv8

The YOLOv8 model was fine-tuned on the annotated dataset for detecting individual flies. Its lightweight architecture and speed made it ideal for real-time mobile deployment. Bounding box annotations were processed to match the model’s input format. The unified detection pipeline extracts features directly from convolutional layers for classification and bounding box regression. YOLOv8 uses multiple detection heads at varying scales, enabling robust detection of objects across sizes [19].

### 5. Detectron2

Used as a robust detection framework with its feature pyramid network (FPN) for multiscale object detection. The model was fine-tuned on the annotated dataset, leveraging pre-trained weights. Feature Extraction Method: Mask RCNN with ResNet-50FPN:Backbone: ResNet-50 with Feature Pyramid Network (FPN). Uses a Region Proposal Network (RPN) to generate candidate regions. Region-based pooling: Regions of Interest (ROIs) are refined through ROIAlign, such as level predictions.[20]

#### Model Training

This study explores the training of five deep learning models for image classification and object detection: ResNet50, InceptionV3, MobileNetV2, YOLOv8, and Detectron2. Each model training process was optimized for the best performance using hyperparameter values considered appropriate for data augmentation and optimization strategies.

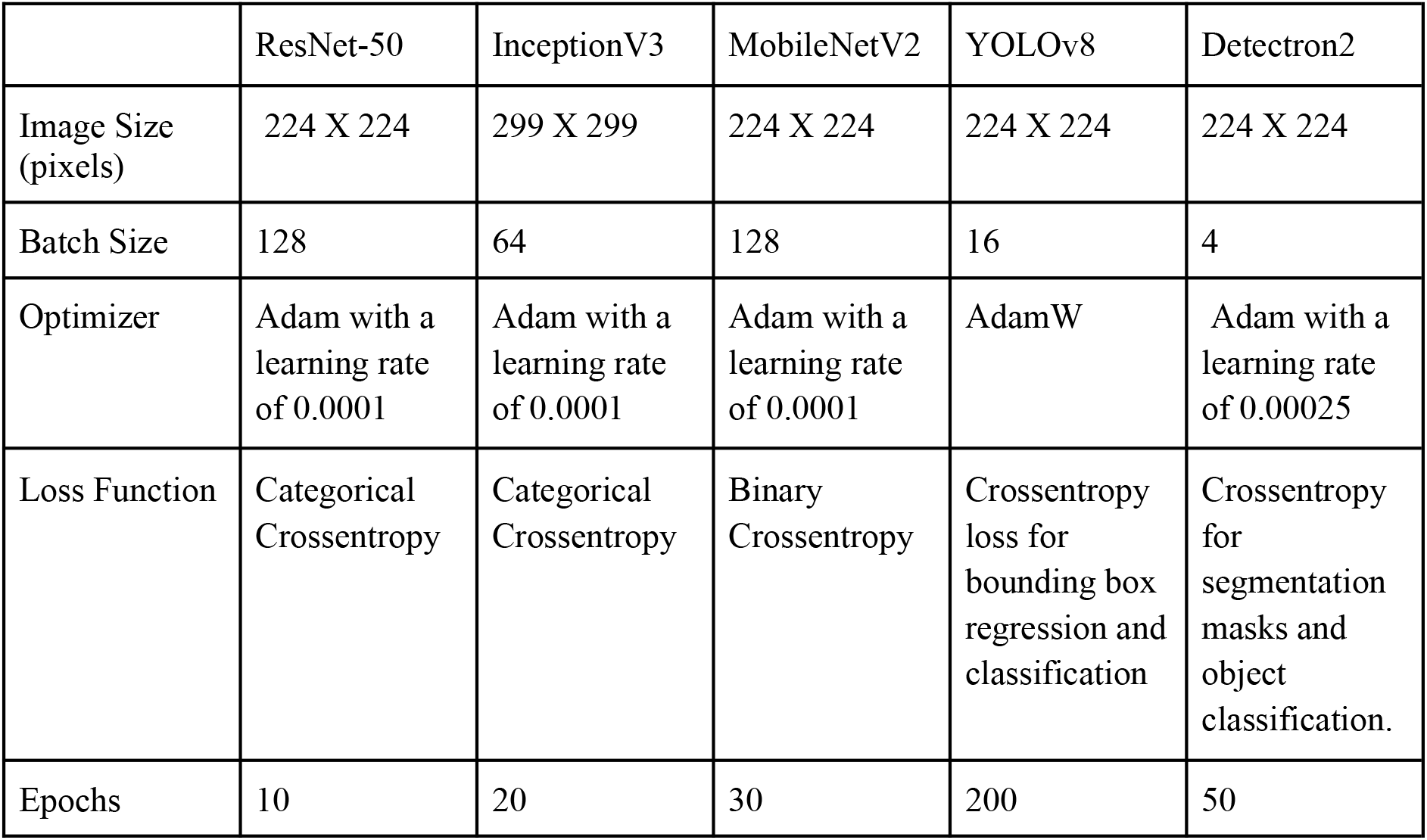

#### Model Performance Metrics

The performance evaluation of the experimental models utilized key metric components based on classification outcomes. True Positives (TP) represent instances where the model accurately identified the positive class, corresponding to the diagonal elements of the confusion matrix. True Negatives (TN) indicate correct negative class predictions, computed as the sum of elements excluding those in the row or column of the class under consideration. False Positives (FP) denote incorrect positive class predictions, while False Negatives (FN) represent missed positive class identifications. These components are instrumental in assessing diagnostic test sensitivity and specificity.The following evaluation metrics were employed in this study:

Accuracy quantifies the model’s overall predictive performance across all data points. While particularly effective for balanced datasets, its utility diminishes for imbalanced class distributions due to potential bias in interpretation.[27]

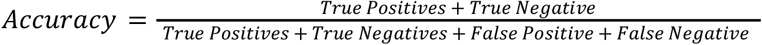

Precision measures the model’s positive predictive value, specifically its ability to accurately identify true positive cases while minimizing false positive predictions. This metric is especially critical in medical diagnostics, where false positives may lead to unnecessary interventions.[26][27]

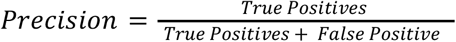

Recall evaluates the model’s sensitivity in detecting positive instances within the dataset. In medical screening applications, recall becomes paramount due to the significant implications of missing positive cases (false negatives).[26][27]

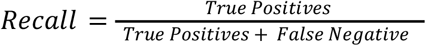

F1-Score provides a harmonized measure of precision and recall, ranging from 0 to 1. A score of 1 indicates perfect prediction across all classes, while 0 signifies complete misclassification. This metric is particularly valuable when addressing non-uniform class distributions.[27]

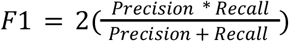

## Result and Discussion

Previous studies utilized various machine learning models for *Drosophila melanogaster* gender classification, their code implementations and applications were not publicly available, making their results difficult to reproduce. This limitation is evident in the following comparison table:

| SNo. | Model Used | Accuracy | F1 score | Precision | Recall | Codes Available & APP |
| --- | --- | --- | --- | --- | --- | --- |
| 1 | AlexNet[21] | 91.25% ± 2.38% | - | - | - | No |
| 2 | SVM[21] | 87.5% ± 1.92% | - | - | - | No |
| 3 | BPNN[21] | 88.75 ± 1.92% | - | - | - | No |
| 4 | YOLOv4-tiny-base[19] | 76.6% | 0.766 | 0.628 | 0.981 | No |
| 5 | YOLOv4-tiny-synt[19] | 81.9% | 0.819 | 0.700 | 0.980 | No |
| 6 | YOLOv4-tiny-synt + mosaic[19] | 83.8% | 0.838 | 0.726 | 0.991 | No |

Performance assessment of models is a necessary step within the entire workflow that seeks to guarantee functionality and trustworthiness. For models, indices like accuracy, precision, recall, and the F1 score were calculated. Of all the models tried and tested, ResNet-50 was attaining a staggering 91% accuracy and MobileNet showed an accuracy of 91% and 63%, respectively.The YOLOv8 mAP model also had the added advantage of being the fastest, with an’mAP’ of 98% (Fig A), making it fit for even real-time usage, especially on mobile devices. This also implies that individual flies could be qualitatively and quantitatively analyzed in images containing many of them and across different scenarios.

| SNo. | Model Used | Accuracy | F1 score | Precision | Recall |
| --- | --- | --- | --- | --- | --- |
| 1 | ResNet-50 | 0.91 | 0.91 | 0.91 | 0.91 |
| 2 | InceptionV3 | 0.78 | 0.78 | 0.78 | 0.78 |
| 3 | MobileNet | 0.62 | 0.62 | 0.62 | 0.62 |
| 4 | Detectron2 | 0.78 | 0.77 | 0.76 | 0.80 |
| 5 | YOLOv8 | **0.98** | 0.98 | 0.98 | 0.98 |

### Mobile Deployment

Streamlit was first used to create a simple web-based tool showing the accuracy and capability of the underlying models. We modified the tool into an Android mobile app to enhance its availability. By allowing researchers and professionals to use the tool in the laboratory, this mobile adaptation greatly increases its convenience and range of uses. The mobile platform included all five models, therefore allowing users to choose the one most pertinent to their situation. Within the same frame, the object detection model may separate and identify many flies—male or female. Users may either take fresh photographs straight via the program interface or upload already existing ones (see Figure 5 for the approach).First, the program was running on the organization local server. The deployment method has been simplified for more general usage, so that anyone may copy the GitHub repository and following the given Steps, therefore enabling simple installation on other servers. This simplicity of use guarantees that continuous improvements to the program may be easily communicated.

**Fig 5:**
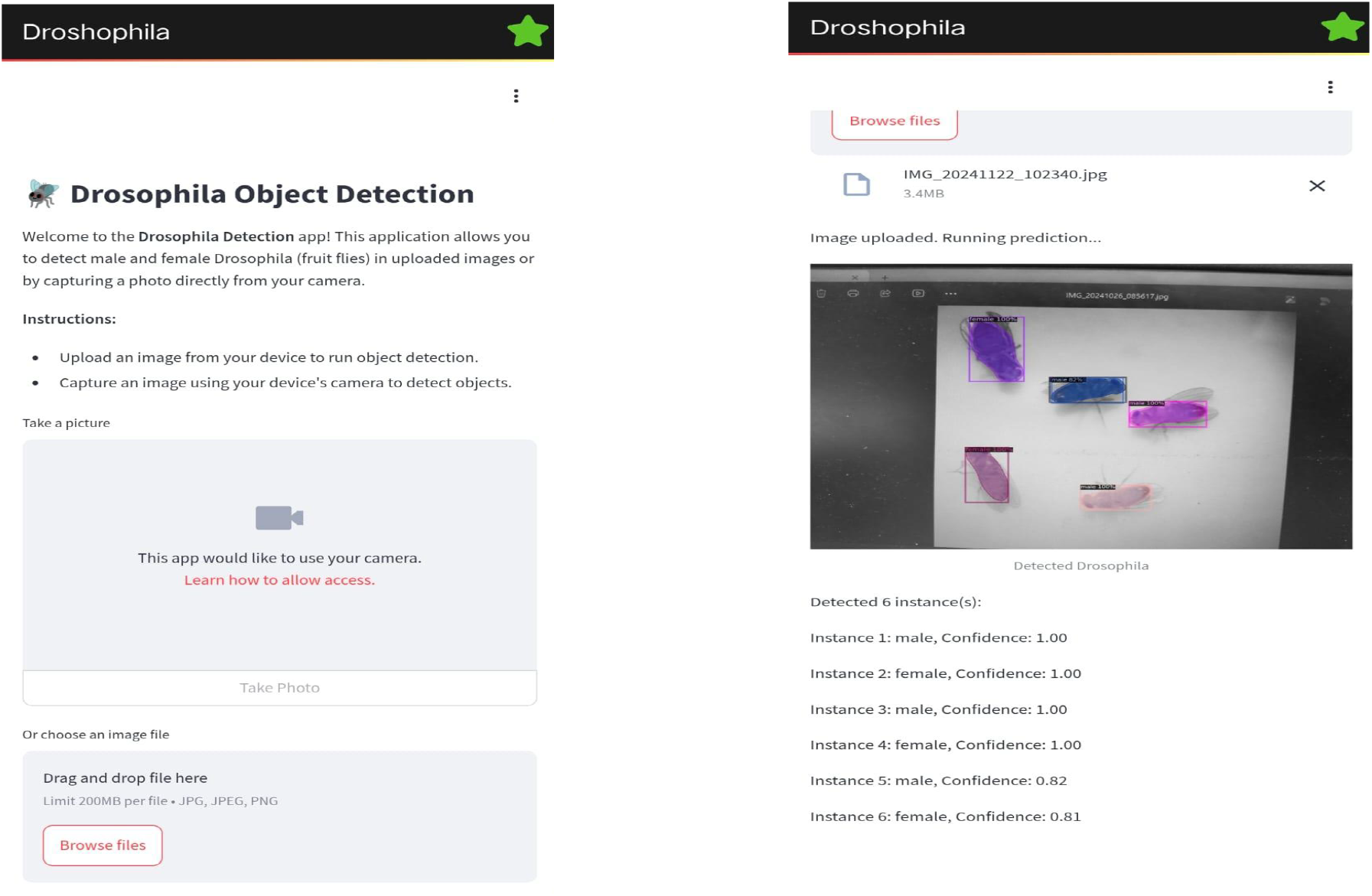
Mobile application interface and object detection

## Conclusion

This work proposes a method for the gender classification of *Drosophila melanogaster* with the use of mobile technology and advanced machine learning algorithms. The research uses advanced object detection models, namely, YOLOv8 and Detectron2, as well as classification architectures such as ResNet-50, InceptionV3, and MobileNet to automate the time-consuming and tedious task of sorting flies by hand. The Oregon-R strain of *Drosophila melanogaster* was imaged using the T60N Series stereo microscope as part of the methodology, followed by dataset creation and preprocessing steps. The classification models, however, employed transfer Learning approaches where the ResNet-50 model used the ImageNet weights, the InceptionV3 relied on its Inception modules, while MobileNet was selected due to its lightweight structure conducive for mobile application. For the purpose of object detection, annotating sites were used as the basis for fine-tuning YOLOv8 and Detectron2. This combination of models embedded within a mobile application showed great potential in aiding real-time gender classification on flies, substantiating the use of deep learning algorithms within biological studies. This method makes it possible to achieve scalable and reproducible and, at the same time, efficient means of classification of the *Drosophila melanogaster* by gender, which in return can greatly improve the rate of work in research focusing on genetics, developmental biology, and disease modeling. The study reported a maximum accuracy of about 98.0% with the application of the YOLOv8 model, which provided proof for the viability of the algorithm assessment framework to the problem of manual classification in terms of time, researcher differences, and the level of skill required to discern forms.

## Code Availability & Supplementary materials

GitHub Link:-https://github.com/jayantrai12/Drosophila_Classification

